# Fetal Bovine and Calf Serums Differ in Their Contents of Endocannabinoids, Unsaturated Fatty Acids, Monoacyl-Glycerols, *N*-Acyl-Ethanolamines, Oxylipins and Cytokines

**DOI:** 10.64898/2025.12.15.694409

**Authors:** Hilal Kalkan, Jean-Philippe C Lavoie, Isabelle Bourdeau-Julien, Frédéric Raymond, Vincenzo Di Marzo, Nicolas Flamand

## Abstract

Cell culture relies heavily on serum supplementation, but serum composition makes it difficult to ensure reproducibility and consistency of experimental results. Consequently, research groups must test multiple batches to ensure the functionality and reproducibility of their models of their models. The commonly used serums are fetal bovine serums (FBS) and calf serums (CS), which have been recognized as crucial in modulating cellular processes. While having been utilized for decades, little information is known about their respective lipid mediator contents. This study explored the presence of several major bioactive lipids involved in the regulation of inflammation, metabolism, differentiation, immune response, neuroprotection, and vascular homeostasis. These included polyunsaturated fatty acids, monoacylglycerols (MAGs), *N*-acyl-ethanolamines (NAEs), and oxylipins. As compared to FBS, CS samples were enriched in most fatty acids except for arachidonic acid. The levels of the endocannabinoids 2-arachidonoyl-glycerol (2-AG) and *N*-arachidonoyl-ethanolamine (AEA) followed the same pattern as arachidonic acid. On the other hand, most lipoxygenase-derived mediators, including, leukotriene B_4_, showed higher abundance in CS. Accordingly, CS serum activated the random migration of human neutrophils to a much greater extent than FBS, an effect attenuated by the BLT_1_ receptor antagonist CP 105,696. These findings highlight how bovine serum lipid composition is a major determinant modulating cellular responses and might thus impact experimental reproducibility. The data presented herein will help key insights beyond conventional cell culture optimization and might represent key features to consider when planning *in cellulo* experiments across numerous fields, also beyond immunology.

## INTRODUCTION

Cell culture are one of the cornerstones of modern biological research, providing highly controlled conditions under which researchers can probe into the detailed functions of cellular machinery, signaling cascades, and physiological responses. Addition of serum in culture media is indispensable as it supplies a cocktail of essential nutrients, lipids, hormones, and growth factors that are critical for maintaining cell viability, supporting proliferation, and facilitating differentiation (1-5). Fetal bovine and calf serums (FBS and CS, respectively) are especially appreciated for their ability to support a wide range of *in cellulo* models (6-8). However, one of the significant drawbacks of serum-containing media is the biochemical variability among different serum types and even batches (5, 9, 10). This inconsistency often leads to undesired variability in experimental outcomes.

Traditionally, lipids have been mainly viewed as structural constituents of cellular membranes or as energy reservoirs. Nowadays, numerous lipid mediators are recognized as highly active signaling molecules affecting almost every aspect of cell function. Fatty acids, monoacyl-glycerols (MAGs), *N*-acyl-ethanolamines (NAEs) and their oxidized metabolites are among the most critical bioactive lipids. Fatty acids are essential in cell metabolism, as both an energy source and precursors for the biosynthesis of several potent lipid mediators, including endocannabinoids, octadecanoids, eicosanoids and docosanoids (11, 12). Most of these mediators are major players in either inflammation, differentiation, vascular & homeostatic responses, as well as repair phenomena, which are typical cell signaling mechanisms responsible for metabolic control. Collectively, these lipid mediators work in concert to execute a wide array of biological functions, underscoring their importance in unraveling the complexities of cellular responses in vitro and in vivo. Importantly, the vast majority of lipid mediators mentioned above are found in all mammals.

Differences in the concentration of numerous bioactive lipids between FBS and CS samples may thus lead to unpredictable changes in cellular responses, especially within research areas related to immunology/inflammation and metabolism in which lipid mediators are heavily involved. Herein, we analyzed the cytokine and lipid mediator content of different serum samples in order to define and highlight the possible differences between FBS and CS, in the hope of helping researchers to select the type of serum they should utilize in their cellular models depending on such information. We also quantitated some bovine cytokines and chemokines, as some cytokines/chemokines were previously shown to activate their ortholog receptors, (13-18) sometimes with comparable efficacy, sometimes with a weaker efficacy compared to human proteins. This study bridges these critical gaps in knowledge by providing an in-depth lipidomic comparison of FBS and CS.

## MATERIAL AND METHODS

### Material

Optima LC-MS Grade water, methanol, acetonitrile, glacial acetic acid, chloroform, Hepes, dimethyl sulfoxide (DMSO) and bovine serum albumin were purchased from ThermoFisher. Hank’s balanced salt solution (HBSS) and the lymphocyte separation medium were from Wisent. Ammonium acetate and dextran were purchased from Millipore Sigma. Internal standards and pure LC-lipid mediators were obtained from Cayman Chemical or synthesized in-house. Anti-CD16 antibody was purchased from Miltenyi Biotec, and 24-well plates with 3 µm pore inserts were from VWR.

### Lipid extraction

Lipids were extracted from FBS and CS samples (Supplementary Table 1) as described in (19) with slight modifications. In brief, serum samples (200 µL) were mixed with 300 µL of HBSS, then denatured with methanol containing a cocktail of deuterated internal standards. The denatured samples were then acidified with acetic acid (final concentration 0.575%). CHCl_3_ (1 ml) was then added to the samples, vortexed for 1 minute, and subsequently centrifuged at 3000 × *g* for 5 minutes. The organic phases were harvested and dried in a rotary evaporator. The dried extracts were dissolved in 60 µl of a 50/50 (v/v) mixture of solvent A (H_2_O containing 1 mM ammonium acetate and 0.05% acetic acid) and solvent B (MeCN: H_2_O, 95:5, v/v, containing 1 mM ammonium acetate and 0.05% acetic acid).

### Lipid mediator analyses by LC-MS/MS

Lipids extracts were injected (40 µl) onto a LC column (Kinetex C8, 150 × 2.1 mm, 2.6 μm, Phenomenex) and eluted at a flow rate of 400 μl/min using a discontinuous gradient of solvent A and B (15–35% B from 0 to 2 min, 35–75% B from 2 to 12 min, 75– 95% B from 12 to 12.1 min and kept at 95% until 17 min). The LC system was interfaced with the electrospray source of a Shimadzu 8050 triple quadrupole mass spectrometer, and mass spectrometric analysis was performed using multiple reaction monitoring with the specific mass transitions listed in Supplementary Table 2 for the compounds and their deuterated homologs or surrogates. Samples were analyzed in a blinded fashion, and only peaks with the same retention time as the purified compound of interest and a signal-to-noise ratio greater than 5 were retained for quantification, as recently recommended (20). In the case of monoacylglycerols (MAGs), the data are presented as 2-MAGs but represent the combined signals from the *sn*-2 and *sn*-1(3) isomers due to acyl migration between the different *sn* positions.

### Cytokine analyses

Cytokine levels in serum samples were assessed by Eve Technologies Corp (Calgary, AB, Canada) by using the Bovine Cytokine/Chemokine Array 15-Plex (BD15). Cytokine/chemokine levels are reported as pg/ml of serum. The cytokines/chemokines analyzed were INFγ, IL-1α, IL-1β, IL-4, IL-6, IL-8, IL-10, IL-17A, IL-36RA, IP-10, MCP-1, MIP-1α, MIP-1β, TNFα, and VEGF-A.

### Isolation of human neutrophils

Experimental protocols were approved by the Research Ethics Committee of the Centre de recherche de l’Institut universitaire de cardiologie et de pneumologie de Québec. Volunteers gave their informed consent. Peripheral blood (160 ml) was collected from healthy volunteers in K_3_EDTA-containing tubes. Neutrophils were next isolated exactly as in (21). Cell viability and purity were ≥ 98% as assessed by trypan blue exclusion and differential staining (hematoxylin/eosin), respectively.

### Migration assays

Transmigration assays were performed using freshly isolated neutrophil suspensions as described in (38345417). In brief, neutrophil suspensions (5 million/ml) were prewarmed at 37 °C in HBSS containing 10 mM Hepes, 1.6 mM CaCl2, and either 5% FBS or 5% CS. Neutrophils were then treated for 15 min with the leukotriene B_4_ receptor antagonist CP 105,696 or vehicle (DMSO). Migration was induced with 100 nM LTB4 or vehicle (DMSO), which were added to the lower chambers of the transmigration apparatus. Neutrophil suspensions (100 µl) were seeded in the upper chambers and were allowed to migrate for 2 h. The contents of the lower chambers were harvested, and neutrophils were counted using a hemacytometer. The serums used for transmigration assays were FBS10 and CS2 for FBS and CS, respectively (Supplementary Table 1).

### Statistical analyses

Statistical analyses were performed using GraphPad (supplementary figures) or R studio software (RStudio 2024.09.1+394, R version 4.4.2). Principal component analysis (PCA) plot was made with the FactoMineR and the factoextra package. Permutational multivariate analysis of variance (PERMANOVA) has been made using Adonis2 function of the package vegan with 100’000 permutations. Hierarchical clustering from the PCA analysis were made using the HCPC function of the hierarchical clustering on FactoMineR package. Heatmaps were drawn using pheatmap package and to allow a better visualization of the overall lipid and bovine cytokines, concentrations were normalized with the preprocess() function of the ‘caret’ package. Zero and NA values were imputed, then all the concentrations were scaled and centered. Volcano plot and boxplots were drawn with ggplot2 package and statistical analysis was done with wilcox.test() function from the stats package. Spearman correlations were calculated using the function cor.test from the stats package, p.values were corrected by FDR using p.adjust() function of the stats package and correlation plot was drawn with corrplot package.

## RESULTS

In order to measure cytokines, we utilized a 15-Plex Discovery Assay (Eve Technologies, Calgary, AB, Canada). All proteins were either detected in FBS, CS, or both. As for the lipid mediators, we screened a total of 99 analytes (Supplementary Table 2), out of which 48 were consistently found in most samples.

To explore the global biochemical divergence between FBS and CS, we performed principal component analysis (PCA) as well as a volcano plot on the complete dataset of quantified lipid mediators and cytokines. The first two principal components accounted for 32.7% and 16.0% of the total variance, respectively. As shown in Figure 1A, the FBS (blue) and CS (red) groups formed clearly separated clusters driven by the first axis, with non-overlapping 95% confidence ellipses. This separation was statistically validated using PERMANOVA (p = 0.001). The top 20 variables contributing to the sample distribution are shown as loadings (dark grey arrows). FBS samples aligned with higher levels of endocannabinoids (AEA, 2-AG) and arachidonic acid (AA). Of note, AA was almost perpendicular to Axis 1 in the PCA plot (Figure 1A). and higher in FBS *vs*. CS samples (p<0,05, Figure 1B and Supplementary figure 1). In contrast, CS samples showed an increase in octadecanoids, eicosanoids, and some docosanoids, notably those derived from the 5-and 15-lipoxygenase pathways (Figure 1). Furthermore, hierarchical clustering of both lipid mediators and bovine proteins revealed distinct signatures associated with each type of serums, supporting the concept that CS and FBS greatly differ in their content of both lipid mediators and cytokines (Figure 1C). Figures 2 and 3 highlight the lipid mediators that were significantly different between FBS and CS. When we look at the different lipids, CS were enriched in OA, LA, and the n-3 polyunsaturated fatty acids SDA and EPA (Figures 1B and 3). Not surprisingly, monoacylglycerols followed the same trend as polyunsaturated fatty acids (Figure 3,4 and Supplementary Figure 1). In addition, and despite that AA levels are lower in CS samples, many 5- and 15-lipoxygenase metabolites were increased in CS samples vs. FBS samples, along with the cyclooxygenase metabolites TXB_2_ and 12-HHTrE (Figure 3). While the levels of PGF_2α_ were comparable between FBS and CS samples (Supplementary figure 2), the analysis of other cyclooxygenase metabolites (PGD_2_ and PGE_2_ for instance), could not be achieved accurately due to LC-MS traces not being in line with the recent technical recommendation for the analysis of lipid mediators (20). A certain level of variability was also observed within each serum type, indicating potential for batch effects associated with serum provenance and lots (Figure 1C and Supplementary Table 1).

**FIGURE 1.**
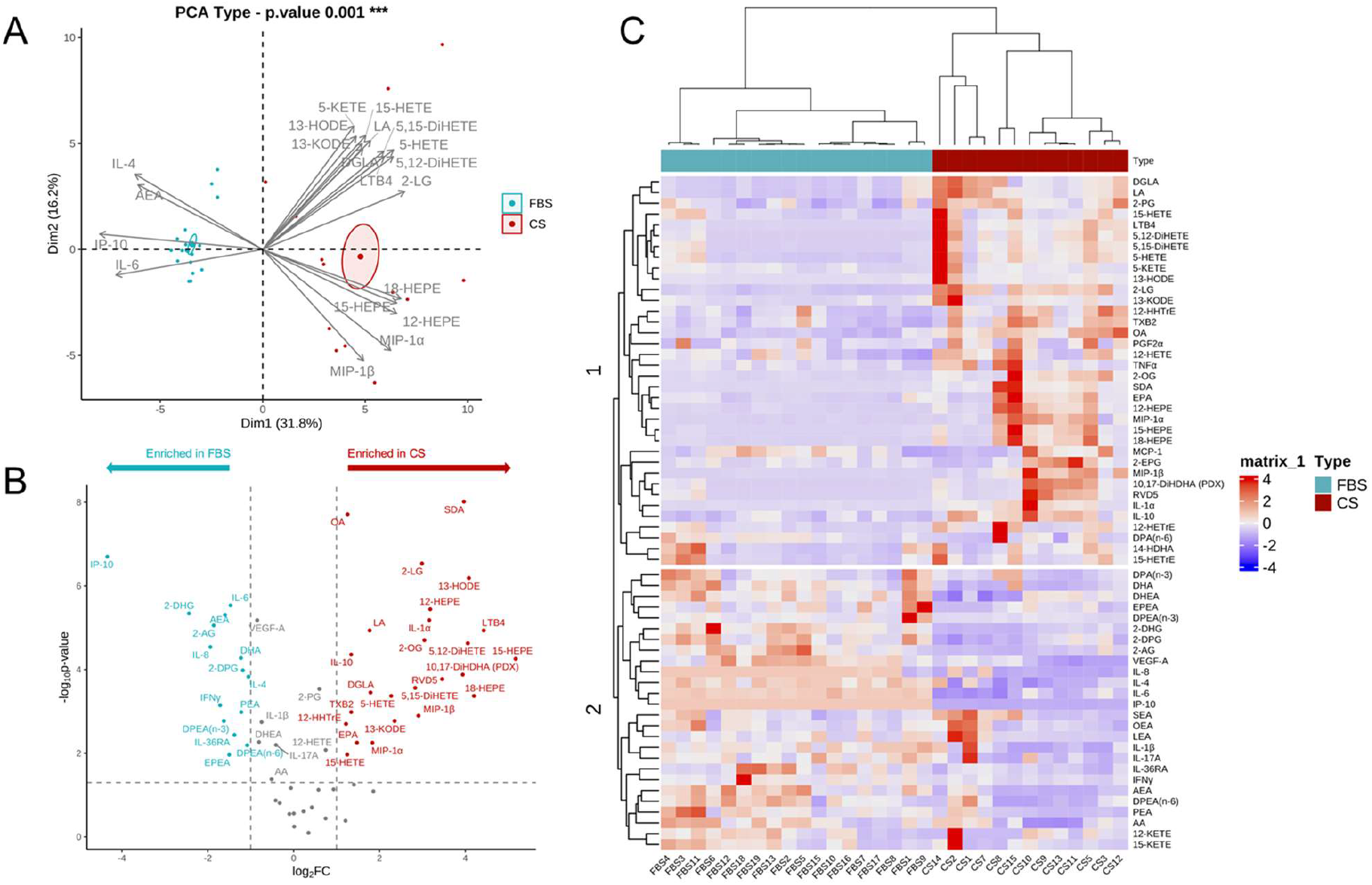
Statistical comparison of FBS and CS samples contents in lipid mediators and cytokines. **A)** PCA of lipid mediators and bovine cytokines in FBS and CS samples. **B) Volcano plot showing the differential content of lipid mediators and bovine cytokines.** The x-axis represents the log2 of the fold change between FBS and CS, and the y-axis represents the –log10 of the *p* value calculated by Wilcoxon rank-sum test. Blue dots indicate metabolites that are significantly enriched in FBS *vs*. CS (p < 0.05, log_2_FC < −1). Red dots indicate metabolite significantly enriched in CS *vs*. FBS (p < 0.05, log_2_FC > 1), blue points indicate, and gray points represent non-significant metabolites. **C) Hierarchical clustering on principal components of lipid mediators and bovine cytokines**. Heatmap representing normalized concentration of lipids and bovine cytokines of CS and FBS media samples. To allow a better visualization of the overall lipid and bovine cytokines, concentrations were normalized with the preprocess() function of the ‘caret’ package. Zero and NA values were imputed, then all the concentrations were scaled and centered.

**FIGURE 2.**
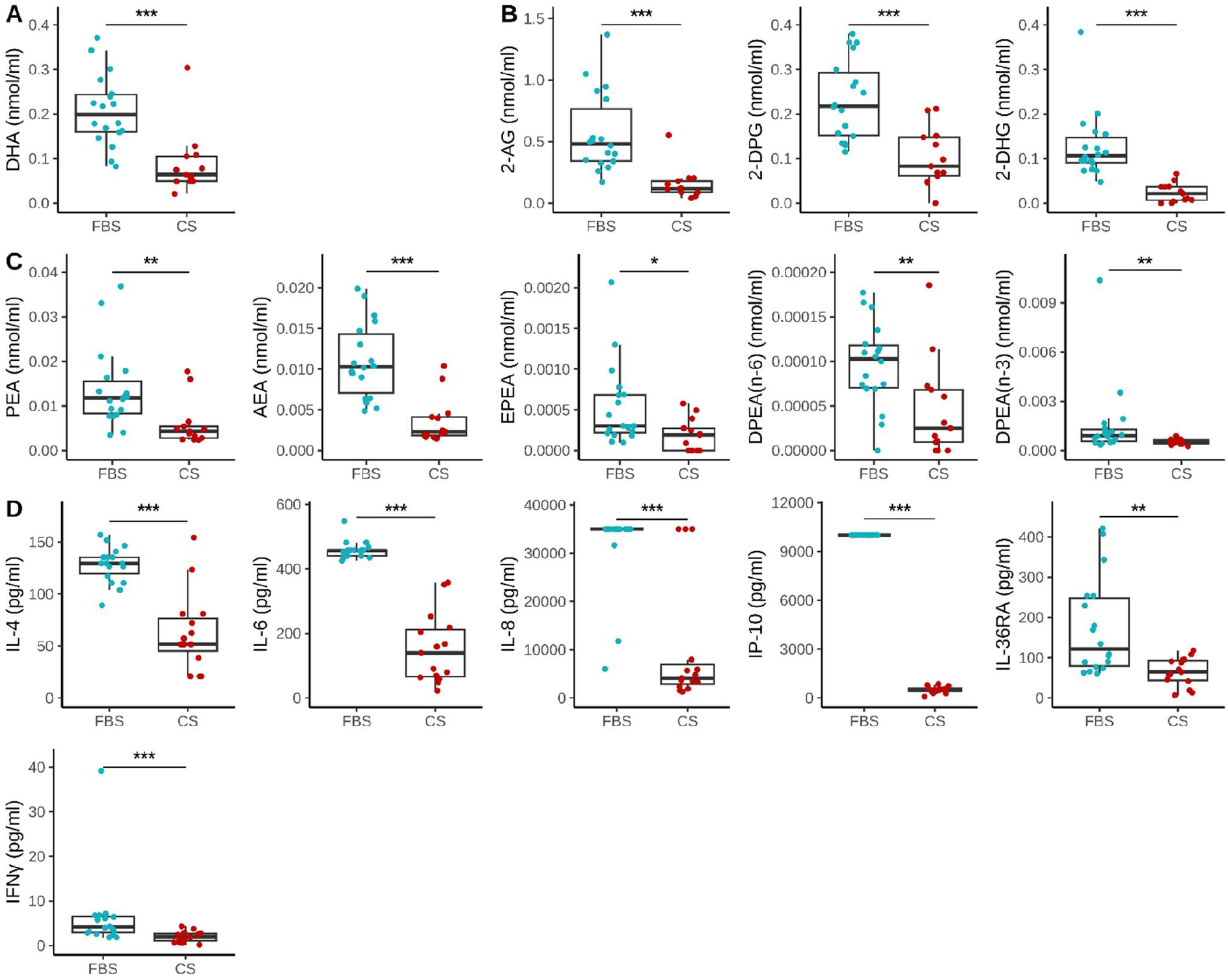
Metabolites significantly enriched in FBS compared to CS (log_2_FC > 1). **A)** Fatty acids; **B)** Monoacylglycerols; **C)** *N*-acyl-ethanolamines; **D)** Cytokines/chemokines. Non-parametric Wilcoxon rank-sum test was performed as most of the metabolites didn’t follow a normal distribution. Significance was set at *p*<0.05 (*), *p*<0.01 (**), and *p*<0.001 (***).

**FIGURE 3.**
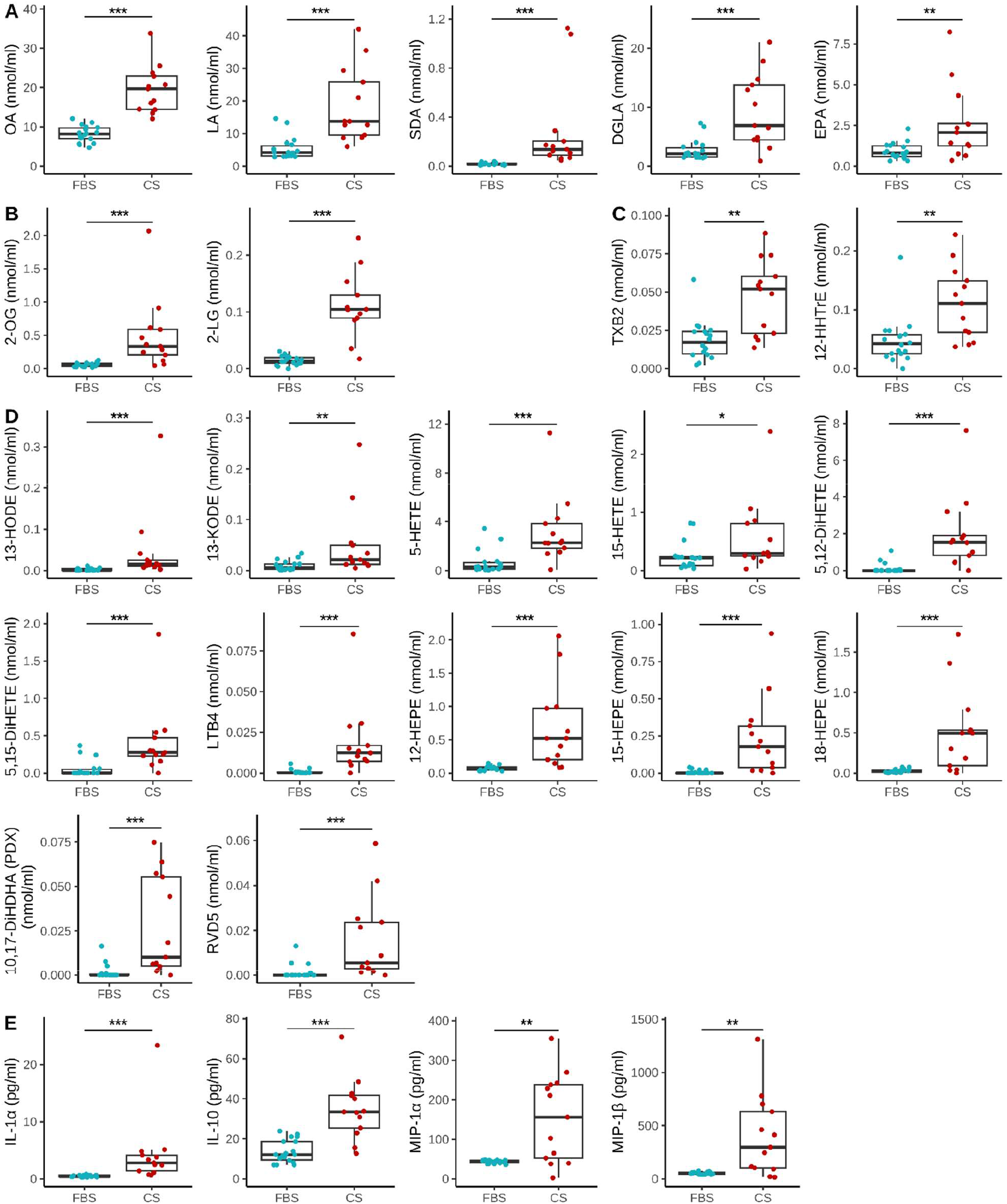
Metabolites significantly enriched in FBS compared to CS (log_2_FC < −1). **A)** Fatty acids; **B)** Monoacylglycerols; **C)** Cyclooxygenase metabolites; **D)** Lipoxygenase metabolites; **E)** Cytokines/chemokines. Non-parametric Wilcoxon rank-sum test was performed as most of the metabolites didn’t follow a normal distribution. Significance was set at *p*<0.05 (*), *p*<0.01 (**), and *p*<0.001 (***).

**FIGURE 4.**
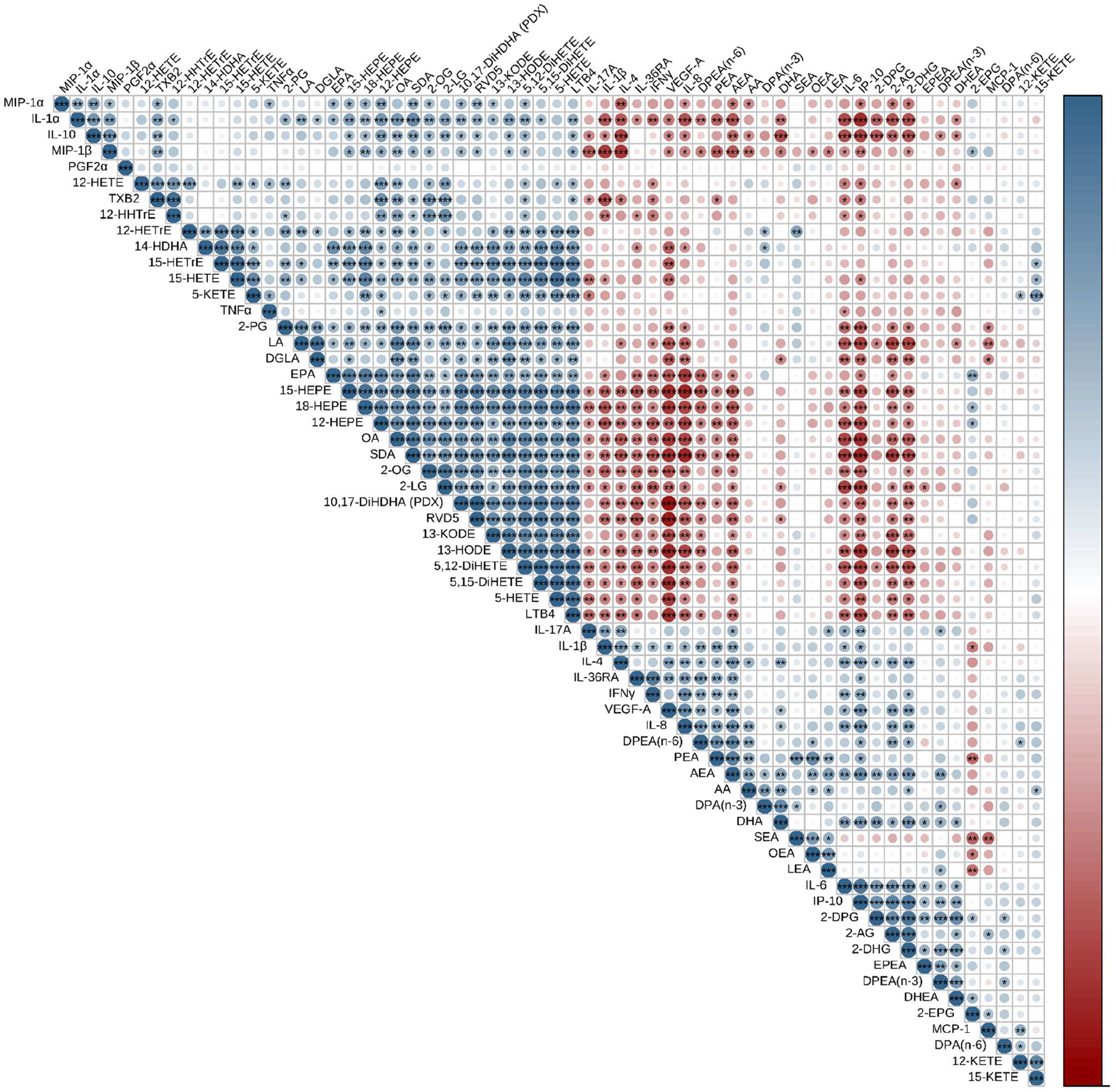
Correlations between lipids and bovine cytokines. Spearman correlations, FDR corrected. Significance was set at p<0.05 (*), p<0.01 (**) and p<0.001 (***).

In light of the higher levels of 5-lipoxygenase metabolites found in CS (*vs*. FBS), we sought to investigate whether this was relevant to the increased random migration of neutrophils that we consistently observed in our migration assay. Therefore, we tested human neutrophil random (unstimulated) migration in presence and absence of the LTB_4_ receptor antagonist CP 105,696, in the attempt to delineate whether the LTB_4_ content of the two types of serums played a role during the assay. In the experiments performed (n=3), isolated human neutrophils barely migrated when incubated in presence of FBS *vs*. CP 105,696 (random migration of 0.42 ± 0.42% and 0.59 ± 0.29% total cells, respectively (mean ± SD; *p* value = 0.39 (t-test)). In contrast, random migration of neutrophils was much greater, and CP 105,696 partially but significantly prevented this (29.3 ± 10.4% *vs*. 17.9 ± 13.9 % respectively, (mean ± SD, *p* value = 0.461 (t-test)). This finding supports the concept that the lipid mediator content of the different serums can impact cellular functions when assessed *in vitro*, at least when assessing human neutrophil functions involving bovine serum.

Several cytokines and chemokines were also found in different levels in FBS and CS. Indeed, IL-4 and IL-6, as well as the CXC chemokines CXCL8 (IL-8) and CXCL10 (IP-10) were found in greater concentrations in FBS samples. In contrast, CS samples had higher levels of IL-10 and the CC chemokines CCL3 and CCL4 (Figures 1,3,4 and supplementary figures). IL-10, IL-6, CXCL8, and CXCL10, were also key drivers of FBS clustering. Interestingly, most CC chemokines that we analyzed better defined CS rather than FBS. Overall, the data presented in Figures 1-3 confirm that the molecular landscapes of FBS and CS are not defined by isolated markers but by coordinated, multidimensional differences in lipid and cytokine composition.

Finally, we also examined the potential links between lipid mediators and bovine cytokines using Spearman correlations. As expected, correlations between the different lipid mediators and cytokines analyzed in Figures 1-3 (and supplementary figures) were also present. In addition, it outlines that the differences observed in both lipid mediators and cytokines between FBS and CS are evident in these groups, with apparent clustering between both lipid mediators and proteins (Figure 4).

## DISCUSSION

Isolated leukocytes and cultures of other cells are among the foundations of biological research, providing *in cellulo* models in which specific functions, including functional assays and signaling cascades can be probed. The current study was undertaken when we noticed the large difference existing between human neutrophil random migration when performed in presence of FBS or CS. We report that the lipid mediator content of FBS and CS greatly differs. Given the recognized impact of lipid mediators across a plethora of cellular functions, the documented differences we unravel likely impacts numerous aspects of *in cellulo* culture and functional assays. In fact, we provide evidence that **1)** FBS and CS differ in their content in polyunsaturated fatty acids and monoacylglycerols; **2)** CS is characterized by higher levels of lipoxygenase-derived metabolites; **3)** CS is characterized by increased TXB_2_ and 12-HHTrE levels. Furthermore, we also provide evidence that **4)** FBS and CS have different cytokine signatures.

Our understanding is that the production of FBS for research purposes arises from the processing of fetal calves that were discovered during the slaughter of cows for feeding purposes. Fetuses are then rapidly collected and their blood harvested through cardiac puncture. The obtained samples vary in volumes, ranging from 300–500 ml, depending on the gestational stage. In contrast, CS samples are usually obtained by harvesting blood from a major vein from calves that are younger than a year. Given the processing of the different sources of serums and the different nutritional aspects of fetal or very young calves, we expected differences in lipid mediator contents as diet is a recognized factor influencing the levels of these metabolites.

One of the striking differences that we observed was the difference in the levels of the endocannabinoids AEA and 2-AG, as well as AA, between the two serum types. Indeed, FBS samples were characterized by higher levels of endocannabinoids and their precursor, AA. In contrast, CS samples were characterized by an increase in many fatty acids, notably oleic acid (OA), linoleic acid (LA) and stearidonic acid (SDA). Such a decrease in AA and an increase in LA in CS samples are in line with what was documented in 1969 by Ahluwalia and colleagues (22). Endocannabinoid-wise, the lower AA concentrations that we observed in CS samples is in line with the decrease in endocannabinoid levels that we also observed. This discrepancy between FBS and CS samples likely play an important role in cellular models investigating endocannabinoids, notably those involved in inflammation, and metabolic disorders (23-25) and we caution investigators to measure endocannabinoid in the serum that they utilize to minimize batch-to-batch effect in their models.

Interestingly, Ahluwalia and colleagues reported a decrease in oleic acid (OA) levels in CS, while we observed the opposite. This is intriguing and difficult to explain. It might possibly reflect a change in cow diet over the years, with an increase in the intake of soybeans and other oils enriched in OA that have been utilized to increase the dietary intake of this fatty acid, thus boosting milk fat, yield, and digestion. Finally, the levels of stearidonic acid (SDA) levels, which were not documented before, were significantly higher in CS *vs*. FBS samples. This is very interesting, as SDA is gaining interest as a more sustainable source of omega-3 polyunsaturated fatty acids, the most recognized source being *B. arvensis* (ahiflower oil) (26). While some studies indicate that SDA levels can be increased to a certain degree in cattle following the intake of encapsulated Echium Oil (27, 28), the lack of scientific evidence, notably in human breast milk, precludes further speculation on this matter.

Interestingly, we also found increased levels of 5- and 15-lipoxygenase metabolites as well as TXB_2_ and 12-HHTrE in CS *vs*. FBS sample. This likely reflects the important increase in circulating platelets and myeloid leukocytes (which express numerous oxygenases) occurring after birth (29-31) and the stimuli affecting the innate immune system following birth. This hypothesis is strengthened by that fact that octadecanoids, eicosanoids and docosanoids are mostly increased in CS *vs*. FBS, despite the fact that the levels of AA and DHA are lower in CS. In fact, it is possible that the lower AA/DHA levels we observed in CS were due in part to increased AA conversion to eicosanoids/docosanoids. Accordingly, we observed that the basal migration of human neutrophils was significantly higher in presence of CS than FBS. Having in mind that octadecanoids and eicosanoids such as 5-HETE and LTB_4_ are chemotactic factors for neutrophils (32-34), we tested the hypothesis that some lipid mediators (LTB_4_) mediated the increased migration that we observed in presence of CS *vs*. FCS. Indeed, we could confirm that LTB_4_ blockade with the BLT_1_ antagonist CP 105,696 partially but significantly blocked neutrophil migration in CS containing experiments, further supporting the concept that the increase in lipid mediator levels in CS significantly modulates immune cell functions. The incomplete blockade suggests that other actors, most likely additional octadecanoids and eicosanoids (32-34), also play an important role in this effect.

When we assessed the levels of inflammatory cytokines and chemokines, we found that FBS and CS had again completely different profiles. Of the fifteen proteins that were measured, only CCL2/MCP-1 levels were comparable between FBS and CS samples (Supplementary Figure 2). Furthermore, nine proteins were elevated in FBS samples (Figure 2 and Supplementary Figure 1), while five proteins had higher levels in CS (Figure 3 and Supplementary Figure 2). Of note, IL-1α, MIP-1α (CCL3) and MIPβ (CCL4) followed the same trends (higher levels in CS samples), which was somewhat expected given the important role of IL-1α in stimulating the expression and production of the two chemokines. Data regarding cross-reactivity between bovine cytokines/chemokines *vs*. their putative human receptors are not always available (35). However, bovine INF-τ, IL-1β and IL-2 cannot activate human cells even though bovine IL-1β can activate murine cells (36-38). In contrast, bovine IL-4 and IL-10 can modulate human cells to the same extent than their human orthologs (15, 18). Finally, other bovine cytokines documented to activate human cells and/or receptors were not analyzed here, such as oncostatin M, CXCL6, fibroblast growth factor-1, leukemia inhibitory factor, IL-32 and lactoferin (16, 39-41). Altogether, and until we fully understand the impact of bovine cytokines/chemokines on human cells, we recommend, in light of the sharp differences that we observed, to investigate the levels of the different proteins that likely play functional roles in the cellular model investigated.

In conclusion, in light of the significant differences between CS *vs*. FBS samples for most of the mediators quantitated in this study, our advice is to measure what they believe to be key mediators involved in their cellular models, keeping in mind that while this might be a cost-driven issue, it will likely help establishing more robust and reliable cellular models and results.

## Abbreviation used in this paper

2-AG: 2-arachidonoyl-glycerol
AA: arachidonic acid
AEA: *N*-arachidonoyl-ethanolamine
CS: calf serum
FBS: fetal bovine serum
MAG: monoacyl-glycerol
NAE: *N*-acyl-ethanolamines.

---

**Supplementary Table 1.** Serum samples utilized in this study.

| **SERUM NAME** | **PROVIDER** | **CATALOG #** | **LOT NUMBER** |
| --- | --- | --- | --- |
| **FBS1** | Wisent | 080-150 | 115698 |
| **FBS2** | Gibco | 123483 | 263590RP |
| **FBS3** | Wisent | 080-105 | 115710 |
| **FBS4** | Wisent | 080-105 | 115714 |
| **FBS5** | Avantor | 97068-085 | 228K22 |
| **FBS6** | Corning | 35-077-CV | 22023001 |
| **FBS7** | Wisent | 098-150 | 115715 |
| **FBS8** | Wisent | 098-150 | 112733 |
| **FBS9** | Wisent | 080-150 | Unknown - 2019 |
| **FBS10** | Wisent | 080-150 | Unknown - 2021 |
| **FBS11** | Corning | 35-010-CF | 80A12-2 |
| **FBS12** | Gibco | A31607-01 | C3022585RP |
| **FBS13** | Gibco | A47668-01 | U2825973RP |
| **FBS15** | Corning | 35-077-CV | 22023001 |
| **FBS16** | Wisent | 080-105 | 115745 |
| **FBS17** | Corning | 35-077-CF | 15822002 |
| **FBS18** | Wisent | 098-105 | 185747 |
| **FBS19** | Gibco | A31607-01 | 2635906RP |
| **CS1** | Cytiva | SH30541.03 | AG29548253 |
| **CS2** | ATCC | 30-2030 | 80207171 |
| **CS3** | Wisent | 074-150 | 87129 |
| **CS5** | Gibco | 16170-078 | 1847453 |
| **CS7** | ThermoFisher | SH30541.02 | AYK170734 |
| **CS8** | Gibco | 16030-074 | 2657905 |
| **CS8** | Gibco | 26010-066 | 2453222 |
| **CS10** | Gibco | 16010-167 | 2526356 |
| **CS11** | Wisent | 074-110 | 87138 |
| **CS12** | Wisent | 074-110 | 87139 |
| **CS13** | Wisent | 074-110 | 87140 |
| **CS14** | ATCC | 30-2030 | 62262399 |
| **CS15** | Gibco | 16170-086 | 2402160 |

**Supplementary Table 2.** Key analytical parameters utilized to quantitate lipid mediators.

| Analyte | Internal Standard | Ret. Time (min) | ESI | m/z Quantifier | m/z Qualifier | LLOQ (fmol) |
| --- | --- | --- | --- | --- | --- | --- |
| 6-keto PGF<sub>1α</sub> | PGE<sub>2</sub>-d4 | 3,52 | - | 369,30→163,10 | 369,30→245,15 | 500 |
| PGF<sub>3α</sub> | PGE<sub>2</sub>-d4 | 3,99 | - | 351,30→193,30 | 351,30→307,40 | 500 |
| TXB<sub>2</sub>-d4 |  | 4,38 | - | 373,10→173,40 | 373,10→199,35 |  |
| TXB<sub>2</sub> | TXB<sub>2</sub> -d4 | 4,41 | - | 369,20→169,25 | 369,20→195,30 | 50 |
| PGE<sub>3</sub> | PGE<sub>2</sub>-d4 | 4,20 | - | 349,30→269,35 | 349,30→313,35 | 50 |
| PGF<sub>2α</sub> | PGE<sub>2</sub>-d4 | 4,45 | - | 353,40→309,30 | 353,40→192,95 | 50 |
| PGE<sub>1</sub> | PGE<sub>2</sub>-d4 | 4,83 | - | 353,30→273,40 | 353,30→317,30 | 50 |
| PGE<sub>2</sub>-d4 |  | 4,69 | - | 355,20→275,35 | 355,20→319,35 |  |
| PGE<sub>2</sub> | PGE<sub>2</sub>-d4 | 4,69 | - | 351,20→271,15 | 351,20→315,20 | 50 |
| PGD<sub>2</sub> | PGE<sub>2</sub>-d4 | 4,91 | - | 351,30→271,20 | 351,30→233,10 | 50 |
| 1a,1b-dihomo-PGF<sub>2α</sub> | PGE<sub>2</sub>-d4 | 5,49 | - | 381,20→337,40 | 381,20→319,40 | 50 |
| PGB<sub>2</sub>-d4 |  | 6,21 | - | 337,40→179,15 | 337,40→239,15 |  |
| 12-HHTrE | PGE<sub>2</sub>-d4 | 7,95 | - | 279,20→179,25 | 279,20→261,25 | 50 |
| PGF<sub>2α</sub>-EA | PGF<sub>2α</sub>-EA-d4 | 3,61 | + | 380,10→344,30 | 380,10→257,00 | 50 |
| PGF<sub>2α</sub>-EA-d4 |  | 3,61 | + | 384,00→348,20 | 384,00→348,20 |  |
| PGE<sub>2</sub>-EA-d4 |  | 3,69 | + | 382,10→364,35 | - |  |
| PGE<sub>2</sub>-EA | PGE<sub>2</sub>-EA-d4 | 3,70 | + | 378,10→360,25 | 378,10→342,25 | 50 |
| PGD<sub>2</sub>-EA | PGE<sub>2</sub>-EA-d4 | 3,88 | + | 378,10→360,25 | 378,10→342,25 | 50 |
| PGF<sub>2α</sub>-G | PGE<sub>2</sub>-G-d5 | 4,04 | - | 427,20→353,30 | 427,20→91,20 | 500 |
| PGE<sub>2</sub>-G-d5 |  | 4,18 | + | 414,00→396,30 | 414,00→299,25 |  |
| PGE<sub>2</sub>-G | PGE<sub>2</sub>-G-d5 | 4,21 | + | 444,25→391,20 | 444,25→409,30 | 50 |
| PGD<sub>2</sub>-G | PGE<sub>2</sub>-G-d5 | 4,37 | + | 444,25→391,30 | 444,25→409,30 | 50 |
| SDEA | EPEA-d4 | 9,17 | + | 320,20→62,15 | 320,20→91,10 | 50 |
| EPEA | EPEA-d4 | 10,15 | + | 346,20→62,20 | 346,20→91,10 | 50 |
| EPEA-d4 |  | 10,10 | + | 350,20→66,15 | 350,20→91,20 |  |
| LEA | LEA-d4 | 10,94 | + | 324,30→62,05 | 324,30→44,00 | 50 |
| LEA-d4 |  | 10,90 | + | 328,00→66,10 | 328,00→48,10 |  |
| DHEA | DHEA-d4 | 11,11 | + | 372,20→62,20 | 372,20→91,15 | 50 |
| DHEA-d4 |  | 11,06 | + | 376,30→66,00 | 376,30→91,10 |  |
| AEA-d4 |  | 11,08 | + | 352,10→66,20 | 352,10→91,20 |  |
| AEA-d8 |  | 11,04 | + | 356,40→44,15 | 356,40→63,25 |  |
| AEA | AEA-d4 | 11,11 | + | 348,30→62,25 | 348,30→287,30 | 50 |
| PEA | PEA-d4 | 11,44 | + | 300,30→62,10 | 300,30→44,05 | 50 |
| PEA-d4 |  | 11,42 | + | 304,00→62,30 | 304,00→44,10 |  |
| OEA | OEA-d4 | 12,07 | + | 326,00→62,05 | 326,00→44,10 | 50 |
| OEA-d4 |  | 12,03 | + | 330,20→66,10 | 330,20→48,10 |  |
| SEA | SEA-d3 | 13,15 | + | 328,20→62,10 | 328,20→44,00 | 50 |
| SEA-d3 |  | 13,14 | + | 331,10→62,20 | 331,10→44,10 |  |
| 1/2-SDG | 1-AG-d5 | 10,28 | + | 351,20→259,25 | 351,20→161,15 | 1000 |
| 1/2-EPG | 1-AG-d5 | 11,16 | + | 377,10→285,25 | 377,10→303,20 | 500 |
| 1/2-DHG | 1-AG-d5 | 11,98 | + | 403,20→311,20 | 403,20→293,20 | 500 |

| Analyte | ISTD | Ret. Time (min) | ESI | m/z Quantifier | m/z Qualifier | LLOQ (fmol) |
| --- | --- | --- | --- | --- | --- | --- |
| 1/2-LG | 1-AG-d5 | 11,90 | + | 355,10→263,15 | 355,10→245,25 | 5000 |
| 1-AG-d5 |  | 12,08 | + | 384,20→287,25 | 384,20→269,30 |  |
| 1/2-AG | 1-AG-d5 | 11,96 | + | 379,20→287,20 | 379,20→269,25 | 100 |
| 1/2-DPG(n-3) | 1-AG-d5 | 12,46 | + | 405,20→313,30 | 405,20→105,10 | 1000 |
| 1/2-OG | 1-AG-d5 | 12,99 | + | 357,20→265,40 | 357,20→247,40 | 1000 |
| 1/2-PG | 1-AG-d5 | 12,39 | + | 331,00→313,30 | 331,01→57,10 | 1000 |
| SDA | AA-d8 | 11,54 | - | 275,00→231,35 | 275,00→177,30 | 100 |
| DPEA (n-6) | DHEA-d4 | 12,04 | + | 374,20→62,10 | 374,20→91,15 | 50 |
| DPEA (n-3) | DHEA-d4 | 11,47 | + | 374,00→62,20 | 374,00→44,10 | 50 |
| EPA-d5 |  | 12,33 | - | 306,20→262,20 | - |  |
| EPA | EPA-d5 | 12,36 | - | 301,20→257,30 | 301,20→203,25 | 50 |
| DHA-d5 |  | 13,09 | - | 332,20→288,20 | - |  |
| DHA | DHA-d5 | 13,11 | - | 327,50→283,30 | 327,50→229,00 | 500 |
| AA-d8 |  | 13,18 | - | 311,20→267,25 | 311,20→182,85 |  |
| AA | AA-d8 | 13,22 | - | 303,20→259,30 | 303,20→205,15 | 50 |
| DPA-d5 |  | 13,33 | - | 334,40→290,35 | 334,40→236,40 |  |
| DPA (n-3) | DPA-d5 | 13,35 | - | 329,50→285,15 | 329,50→231,10 | 100 |
| DPA (n-6) | DPA-d5 | 13,45 | - | 329,10→285,35 | 329,10→231,30 | 50 |
| LA | AA-d8 | 13,23 | - | 279,20→261,40 | 279,20→58,90 | 50 |
| OA | AA-d8 | 13,58 | - | 281,20→263,45 | 281,20→97,00 | 50 |
| 15-HEPE-EA | 15-HEPE-EA-d4 | 7,01 | + | 344,10→62,20 | - | 50 |
| 15-HEPE-EA-d4 |  | 6,98 | + | 348,30→66,15 | 348,30→283,20 |  |
| 13-HODE-EA-d4 |  | 7,23 | + | 326,30→66,20 | 326,30→48,10 |  |
| 13-HODE-EA | 13-HODE-EA-d4 | 7,25 | + | 322,00→62,20 | 322,00→344,35 | 50 |
| 15-HETE-EA | 15-HEPE-EA-d4 | 7,71 | + | 346,40→62,25 | - | 50 |
| 15-HEPE-G-d5 |  | 7,85 | + | 380,10→283,20 | 380,10→257,05 |  |
| 15-HEPE-G | 15-HEPE-G-d5 | 7,88 | + | 375,20→283,20 | 375,20→131,10 | 50 |
| 12-HEPE-G | 15-HEPE-G-d5 | 8,05 | + | 375,30→283,25 | 375,30→265,30 | 100 |
| 13-HODE-G-d5 |  | 8,21 | + | 358,20→261,20 | 358,20→67,15 |  |
| 13-HODE-G | 13-HODE-G-d5 | 8,25 | + | 353,20→261,20 | - | 50 |
| 17-HDPA-EA | 15-HEPE-EA-d4 | 8,28 | + | 372,10→62,15 | 372,10→130,90 | 50 |
| 15-HETE-G | 15-HEPE-G-d5 | 8,59 | + | 377,25→285,20 | 377,25→267,20 | 50 |
| 17-HDHA-EA | 15-HEPE-EA-d4 | 8,71 | + | 370,20→62,10 | 370,20→309,25 | 500 |
| 17-HDHA-G | 15-HEPE-G-d5 | 8,76 | + | 401,20→309,35 | 401,20→129,05 | 50 |
| 17-HDPA-G | 15-HEPE-G-d5 | 9,19 | + | 403,10→131,15 | 403,10→311,25 | 100 |
| 18-HEPE | 15-HETE-d8 | 8,63 | - | 317,30→259,40 | 317,30→299,35 | 50 |
| 8,15-DiHETE | 15-HETE-d8 | 6,70 | - | 335,40→317,30 | 335,40→235,25 | 500 |
| 5,15-DiHETE | 15-HETE-d8 | 6,85 | - | 335,20→317,25 | 335,20→173,25 | 500 |
| 5,6-DiHETE | 15-HETE-d8 | 8,28 | - | 335,00→317,35 | 335,00→114,95 | 500 |
| 13-HOTrE | 13-HODE-d4 | 8,61 | - | 292,90→195,25 | 292,90→275,25 | 100 |
| 15-HEPE | 15-HETE-d8 | 8,97 | - | 317,40→219,25 | 317,40→255,30 | 50 |
| 12-HEPE | 15-HETE-d8 | 9,16 | - | 317,30→179,35 | 317,30→255,40 | 50 |
| 13-HODE-d4 |  | 9,34 | - | 299,10→198,15 | 299,10→217,15 |  |

| Analyte | ISTD | Ret. Time (min) | ESI | m/z Quantifier | m/z Qualifier | LLOQ<sup>a</sup> (fmol) |
| --- | --- | --- | --- | --- | --- | --- |
| 12-HETE-d8 |  | 10,02 | - | 327,20→184,20 | 327,20→309,30 |  |
| 12-HETE | 12-HETE-d8 | 10,09 | - | 319,10→179,25 | 319,10→301,30 | 50 |
| 9-HODE | 13-HODE-d4 | 9,38 | - | 295,30→171,35 | 295,30→277,35 | 50 |
| 13-HODE | 13-HODE-d4 | 9,39 | - | 295,50→195,30 | 295,50→277,30 | 50 |
| 15-HETE-d8 |  | 9,64 | - | 327,20→226,20 | - |  |
| 15-HETE | 15-HETE-d8 | 6,73 | - | 319,40→219,30 | 319,40→301,25 | 50 |
| 13-KODE | 5-KETE-d7 | 9,76 | + | 295,20→277,30 | 295,20→179,25 | 50 |
| 15-KETE | 5-KETE-d7 | 10,07 | - | 317,20→113,00 | 317,20→273,35 | 50 |
| 11-HETE | 15-HETE-d8 | 9,92 | - | 319,30→167,35 | 319,30→149,20 | 50 |
| 8-HETE | 15-HETE-d8 | 9,99 | - | 319,30→155,25 | 319,30→301,35 | 50 |
| 17-HDHA | 15-HETE-d8 | 9,90 | - | 343,50→281,30 | 343,50→201,30 | 50 |
| 14-HDHA | 15-HETE-d8 | 10,06 | - | 343,50→281,30 | 343,50→161,20 | 50 |
| 7-HDHA | 15-HETE-d8 | 10,06 | - | 343,15→281,20 | 343,15→141,05 | 50 |
| 4-HDHA | 15-HETE-d8 | 10,43 | - | 343,15→281,15 | 343,15→101,00 | 100 |
| 12-KETE | 5-KETE-d7 | 10,39 | - | 317,10→273,35 | 317,10→153,20 | 100 |
| 5-HETE | 5-HETE-d8 | 10,15 | - | 319,30→115,05 | - |  |
| 5-HETE-d8 |  | 10,07 | - | 326,99→309,35 | 326,99→116,00 |  |
| 17-HDPA | 15-HETE-d8 | 10,34 | - | 345,20→327,10 | 345,20→247,35 | 500 |
| 17-oxo-DHA | 15-HETE-d8 | 10,27 | - | 341,20→297,20 | 341,20→111,05 | 1000 |
| 15-HETrE | 15-HETE-d8 | 10,27 | - | 321,50→303,40 | 321,50→221,30 | 100 |
| 12-HETrE | 15-HETE-d8 | 10,51 | - | 321,15→181,10 | 321,15→303,20 | 100 |
| 5-KETE | 5-KETE-d7 | 10,85 | - | 317,49→203,20 | 317,49→273,35 | 100 |
| 5-KETE-d7 |  | 10,78 | + | 326,20→194,20 | 326,20→308,30 |  |
| RVE1 | RVD2-d5 | 3,63 | - | 349,30→195,20 | 349,30→107,10 | 500 |
| RVD2-d5 |  | 4,93 | - | 380,40→175,20 | - |  |
| RVD3 | RVD2-d5 | 4,74 | - | 375,40→147,20 | 375,40→137,20 | 100 |
| RVD2 | RVD2-d5 | 4,95 | - | 375,40→175,20 | 375,40→217,15 | 1000 |
| LXA<sub>4</sub> | LTB<sub>4</sub>-d4 | 5,21 | - | 351,50→115,05 | 351,50→217,15 | 500 |
| RVD1 | RVD2-d5 | 5,31 | - | 375,40→141,10 | 375,40→217,15 | 100 |
| RVD4 | RVD2-d5 | 5,77 | - | 375,40→101,05 | 375,40→357,30 | 50 |
| 5,15-DiHEPE (RVE4) | RVD2-d5 | 6,15 | - | 333,20→315,25 | 333,20→271,30 | 100 |
| 7,14-DiHDHA (Maresin 1) | RVD2-d5 | 6,90 | - | 359,40→177,25 | 359,40→93,15 | 500 |
| 10,17-DiHDHA (PDX) | RVD2-d5 | 6,99 | - | 359,30→153,15 | 359,30→206,25 | 100 |
| RVD5 | RVD2-d5 | 7,00 | - | 359,40→199,25 | 359,40→279,35 | 50 |
| 13,14-DiHDHA (Maresin 2) | RVD2-d5 | 7,66 | - | 359,40→221,05 | 359,40→232,25 | 100 |
| 20-COOH-LTB<sub>4</sub> | LTB<sub>4</sub>-d4 | 3,59 | - | 365,00→347,25 | 365,00→195,20 | 500 |
| 20-OH-LTB<sub>4</sub> | LTB<sub>4</sub>-d4 | 3,71 | - | 351,10→195,15 | 351,10→333,25 | 100 |
| LTB<sub>5</sub> | LTB<sub>4</sub>-d4 | 6,12 | - | 333,40→195,25 | 333,40→315,10 | 100 |
| LTB<sub>4</sub>-d4 |  | 6,92 | - | 339,30→197,20 | 339,30→321,40 |  |
| 5,12-DiHETE | LTB<sub>4</sub>-d4 | 7,23 | - | 334,99→195,10 | 334,99→317,15 | 50 |
| LTB<sub>4</sub> | LTB<sub>4</sub>-d4 | 6,94 | - | 335,30→195,25 | 335,30→317,30 | 50 |
| 12-oxo-LTB<sub>4</sub> | LTB<sub>4</sub>-d4 | 7,51 | - | 333,40→179,20 | 333,40→128,90 | 50 |
| LTB<sub>3</sub> | LTB<sub>4</sub>-d4 | 7,74 | - | 337,40→195,20 | 337,40→319,20 | 50 |

| Analyte | ISTD | Ret. Time (min) | ESI | *m/z* Quantifier | *m/z* Qualifier | LLOQ<sup>a</sup> (fmol) |
| --- | --- | --- | --- | --- | --- | --- |
| DGLA | AA-d8 | 13,43 | - | 305,05→261,10 | 305,05→287,05 | 500 |
| PGB<sub>2</sub> | PGE<sub>2</sub>-d4 | 6,23 | - | 333,30→174,85 | 333,30→315,00 | 50 |

<sup>a</sup>Lowest limit of quantification representing the lowest concentration of our calibration curve for which the signal-to-noise ratio was greater than 5.

**SUPPLEMENTARY FIGURE 1.**
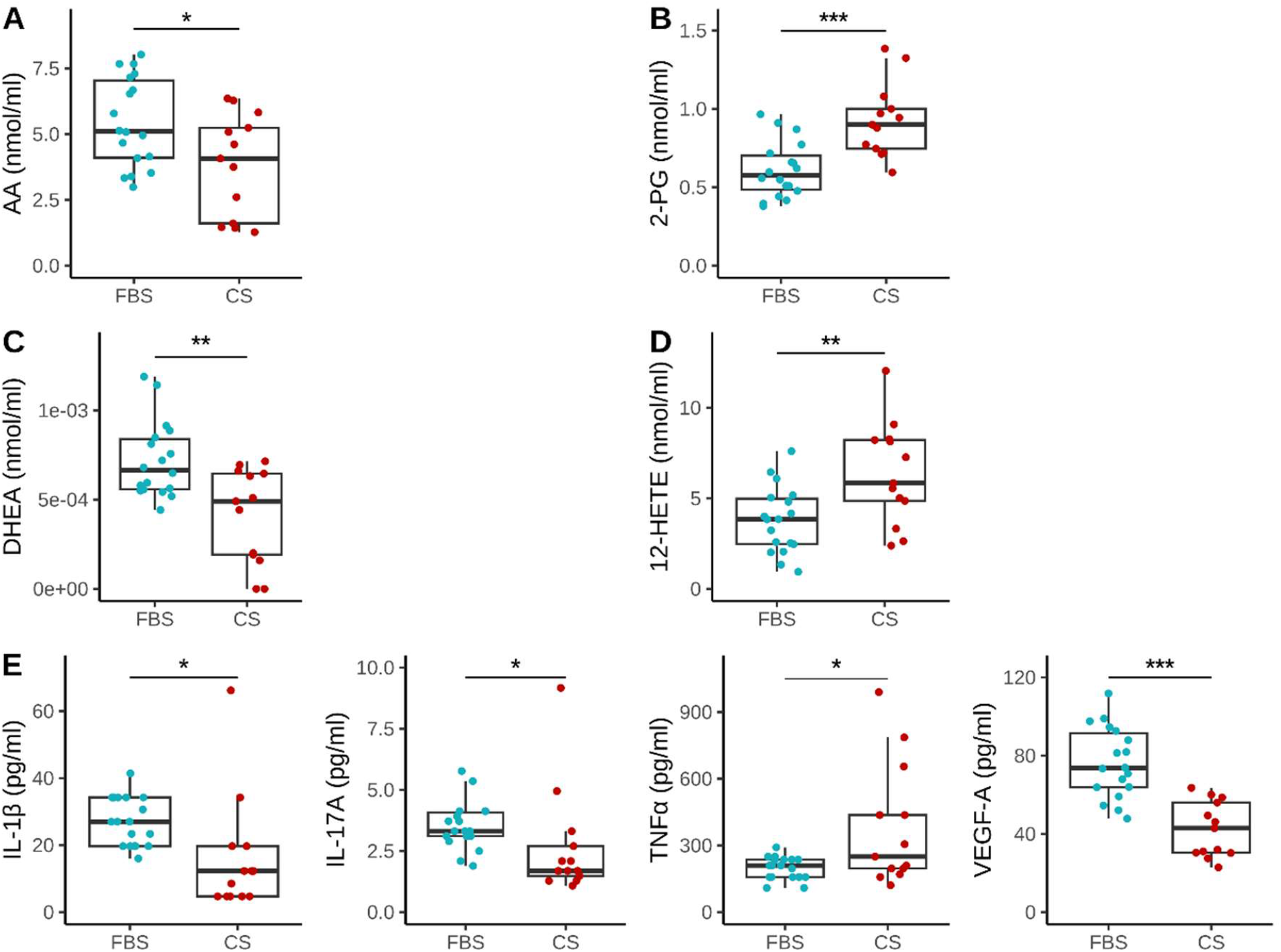
Metabolites significantly different between FBS and CS with a log_2_FC between –1 and 1. **A)** AA; **B)** 2-PG; **C)** DHEA; **D)** 12-HETE; and **E)** Cytokines/chemokines. **A-E)** Non-parametric Wilcoxon rank-sum test was performed as most of the metabolites didn’t follow a normal distribution. Significance was set at *p*<0.05 (*), *p*<0.01 (**), and *p*<0.001 (***).

**SUPPLEMENTARY FIGURE 2.**
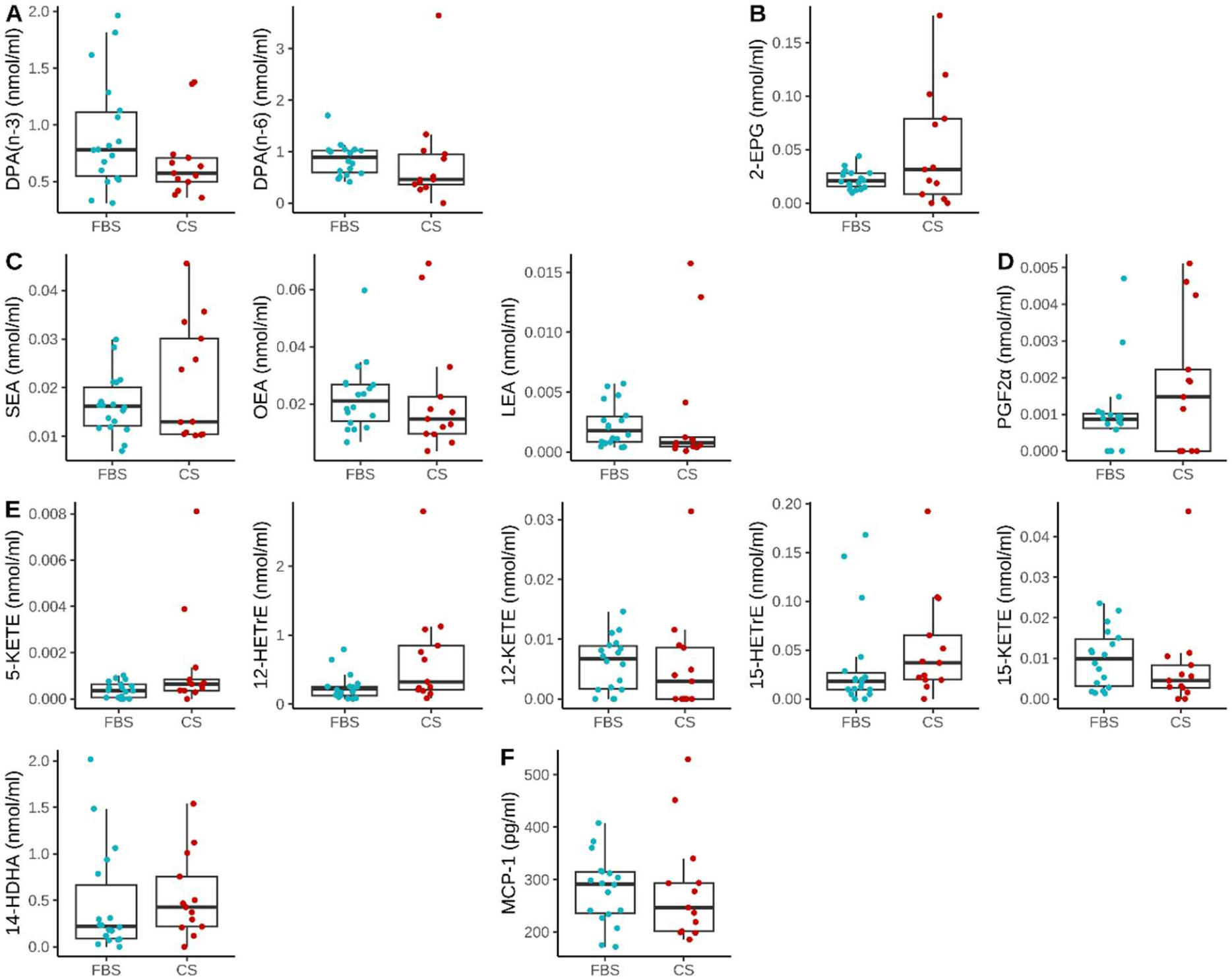
Metabolites that were not significantly different between FBS and CS. **A)** Fatty Acids, **B)** 2-EPG, **C)** NAEs, **D)** PGF_2α_, **E)** lipoxygenase metabolites, **F)** MCP-1 (CCL2). **A-F)** Non-parametric Wilcoxon rank-sum test was performed as most of the metabolites didn’t follow a normal distribution. Significance was set at *p*<0.05 (*).

